# *CDKN1A-RAB44* transcript fusion and activation in cancers

**DOI:** 10.1101/111856

**Authors:** Han Sun, Michelle Nguyen, William Mueller, Zhuanfen Cheng, Hong Zeng, Chenchen Zhu, Jingyan Wu, Kevin Roy, Petra Jakob, Raeka Aiyar, Wu Wei, Lars M. Steinmetz

## Introduction

Splicing contributes to gene regulation and protein diversity, while abnormal splicing underlies both hereditary diseases and cancers^1, 2^. Various mutations that disrupt splicing factors^3, 4^, exonic or intronic splicing enhancers or silencers^5^, as well as splice sites^6^, could be responsible for abnormal splicing. Characterization of abnormal splicing events is not only helpful for understanding the molecular processes linking mutations to disease phenotypes, but also provides promising targets for targeted therapies^7–9^. In addition, CRISPR/Cas9 editing could be benefited once more attention is given to potential abnormal splicing outcomes other than off-target effects at the DNA level^10^. Although large-scale multiplexed genome editing has been demonstrated in yeast^11^, and has also been attempted for particular exons or genes in other eukaryotic cells to achieve saturation^12, 13^, in practice it is much more difficult to measure splicing consequences with genome-wide saturation editing in human cells. Instead, massive somatic mutations accumulated in cancer cohorts provide invaluable opportunities to study somatic mutation-associated splicing events.

Abnormal splicing is not necessarily limited to single genes. Transcript fusion is a special form of abnormal splicing that connects two or more genes due to splicing on a transcriptional level (rather than chromosomal translocations such as *BCR-ABL* in chronic myeloid leukemia). It could arise from conventional splicing on read-through transcripts when the two genes are next to each other and on the same strand, or from trans-splicing when two genes are on different chromosomes, strands or far away—a few cases had been reported^14, 15^. However, it was found that these fusions not only occurred in tumors but also in normal tissues; there was limited investigation regarding how the fusion could happen, whether it be due to mutations or not, and what the downstream perturbations were.

Here, in an effort to characterize somatic mutation-associated abnormal splicing (especially in its simplest form, exon skipping events), we identified over one hundred such events in various tumors, including those in *MET*, *PTEN* and *TP53*. Surprisingly, we detected a recurrent, but previously undescribed, tumor-specific transcript fusion event between the cyclin-dependent kinase inhibitor *CDKN1A* and the *RAS* oncogene family gene *RAB44*. By creating genome-edited cell lines, we demonstrate a causal relationship between splice-site mutations in *CDKN1A* and the fusion to the *RAB44* transcript. We further provide evidence that the fusion arises from a readthrough transcript that escapes exosome-mediated degradation when the splice-site mutation occurred, and we show that the presence of the fusion transcript correlates with *TP53* inactivation and *CDK* activation. The strong tissue specificity of *RAB44* and the relatively high prevalence of this transcript fusion in multiple types of cancers warrants further study which could inform subclassifications of these cancers and the development of targeted therapies.

## Results

### Identification of somatic mutation-associated exon skipping events in cancers

Whole exome sequencing and RNA-Seq data of tumors and matched tumor adjacent controls were collected from 779 patients from TCGA and three other published studies^16–18^, as illustrated in Figure 1A. Taking advantage of Bayesian hierarchical modeling (Supplementary Figure 1), we identified 123 tumor specific exon skipping events associated with somatic mutations in the same exon (Supplementary Table 1). Most frequent were events associated with mutations in splice sites (Figure 1B). In silico prediction of splice site activity^19^ as well as the splice site motif conservation score^20^, before and after mutations, demonstrate the overall accuracy of our identification (Figure 1C and Supplementary Figure 2). In addition, we recaptured a well-known skipping event of exon 14 in a hepatocyte growth factor receptor, *MET*, in two lung cancer patients (Figure 1D). This skipping event (which was suspected to disrupt ubiquitin-mediated degradation and contribute to the oncogenic activation) is an attractive target for small molecule tyrosine kinase inhibitors (TKIs) that are undergoing clinical trials^8^. Besides, we identified exon-skipping events in *MLH1* (a DNA mismatch repair gene), and *ATP6AP2* (a renin receptor gene) as shown in Figure 1E. The same events were previously reported to be causative in hereditary non-polyposis colorectal cancer and Parkinson’s disease respectively^21, 22^, however it remains to be explored whether they are pathogenic to the small-cell lung cancer patient and the head and neck cancer patient here.

**Figure 1.**
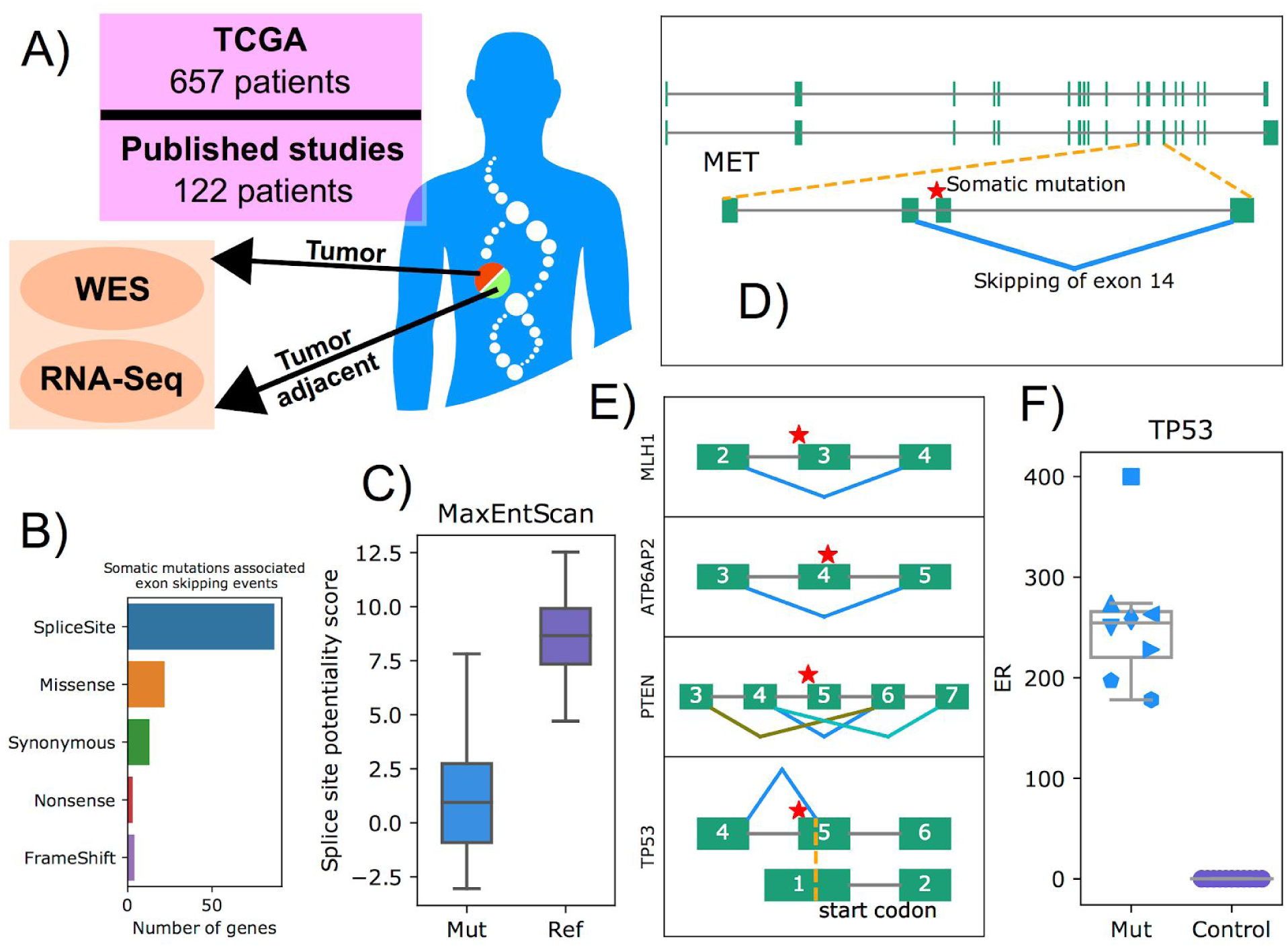
Identification of exon skipping events which associated with somatic mutations. **(A)** Simple project design that both tumors and their matched tumor adjacents data were collected from 779 patidents of TCGA project and three published studies, including both whole exome sequencing (WES) and RNA-Seq. **(B)** Different types of exon skipping events identified. Splice site has the highest annotation priority when a mutation belongs to multiple categories. **(C)** Prediction of splice site activity on the sequences around the splice site, with or without mutations, using MaxEntScan. **(D)** Identification of a well-known exon skipping event in *MET* gene in lung cancer patients. **(E)** Additional interesting somatic mutation-associated exon skipping events. A splice site mutation-associated skipping of exon 3 in MLH1 and a missense mutation, 40bp away from the splice site, associated skipping of exon 4 in *ATP6AP2*. Three different skipping events of exon 5 in *PTEN* in breast cancers. There were reads joining 3rd and 6th exon, 4th and 7th exon as well as 4th and 6th exon. The skipping event of *TP53* utilizes an inner site which is exactly a start codon, indicated by the yellow dot line, according to another isoform. **(F)** In CCLE project, there were 8 cell lines carrying mutations around the splice site of exon 5 in *TP53*. All these cell lines present such skipping events on utilizing the start codon site. Y-axis is the number of reads supporting this skipping event (exclusive reads, ER). Legends for these cell lines are in Supplementary Figure 3A.

We noted the presence of somatic mutations, along with their associated exon-skipping events in *PTEN* and *TP53*. It is interesting that three different events were observed for *PTEN* regarding the skipping of exon 5 (Figure 1E), which can be verified in a leukemia T-cell line (PF-382). For *TP53*, the skipping event is atypical given that it involves utilizing an inner start codon for another isoform (Figure 1E). On the protein level, this could lead to an in-frame deletion of 7aa that is within the transcription factor DNA-binding domain, assuming there is not switching of start codon. Independent evidence from CCLE project shows eight splice mutated cancer cell lines all carry such skipping event (Figure 1F).

### Transcript fusion between *CDKN1A* and *RAB44* activated *RAB44* expression

In addition to the exon skipping events within single genes mentioned above, we observed a novel transcript fusion, associating with a splice site somatic mutation, between cyclin-dependent kinase inhibitor *CDKN1A* and a *RAS* oncogene family member *RAB44* in tumors. The second exon of *RAB44* was joined directly to the first exon of *CDKN1A*, accompanied by the skipping events of exon 2 within *CDKN1A* (Figure 2A). As the start codons of both these two genes located in their second exon, this kind of transcript fusion did not change protein sequence of *RAB44* at all. However, expression of *RAB44* was activated clearly, together with the down regulation of *CDKN1A*, as shown in Figure 2B and 2C. It is worth noting that *RAB44* is not or very lowly expressed, in various human tissues except blood (Supplementary Figure 3B), which also suggests that these tumor cells might be able to escape easily from the T cell-mediated immune surveillance in the sense that T cells probably can’t recognize the peptides from *RAB44* even they could be presented, as they are the same as those in normal blood cells. Furthermore, all the six cancer patients carrying such splice site mutations, either point mutations or small indels, have higher *RAB44* expression, including 4 from 400 bladder cancers, 1 from 400 stomach cancers and 1 from 100 skin cancers (Figure 2D). We proposed a model to explain these observations that it is the splice site disruptive mutations cause the exon skipping events within *CDKN1A* as well as the transcript fusion between *CDKN1A* and its downstream gene *RAB44*. The fusion is further responsible for the activation of *RAB44* on transcription level as shown in Figure 2E. An alternative model, the activation of RAB44 was due to the down-regulation of CDKN1A rather than the direct effect of the fusion, was much less likely given that we couldn’t see the first exon of RAB44 in the RNA-Seq data of the fusion tumors and that we couldn’t see negative correlation on expression between these two genes in general (Supplementary Figure 4).

**Figure 2.**
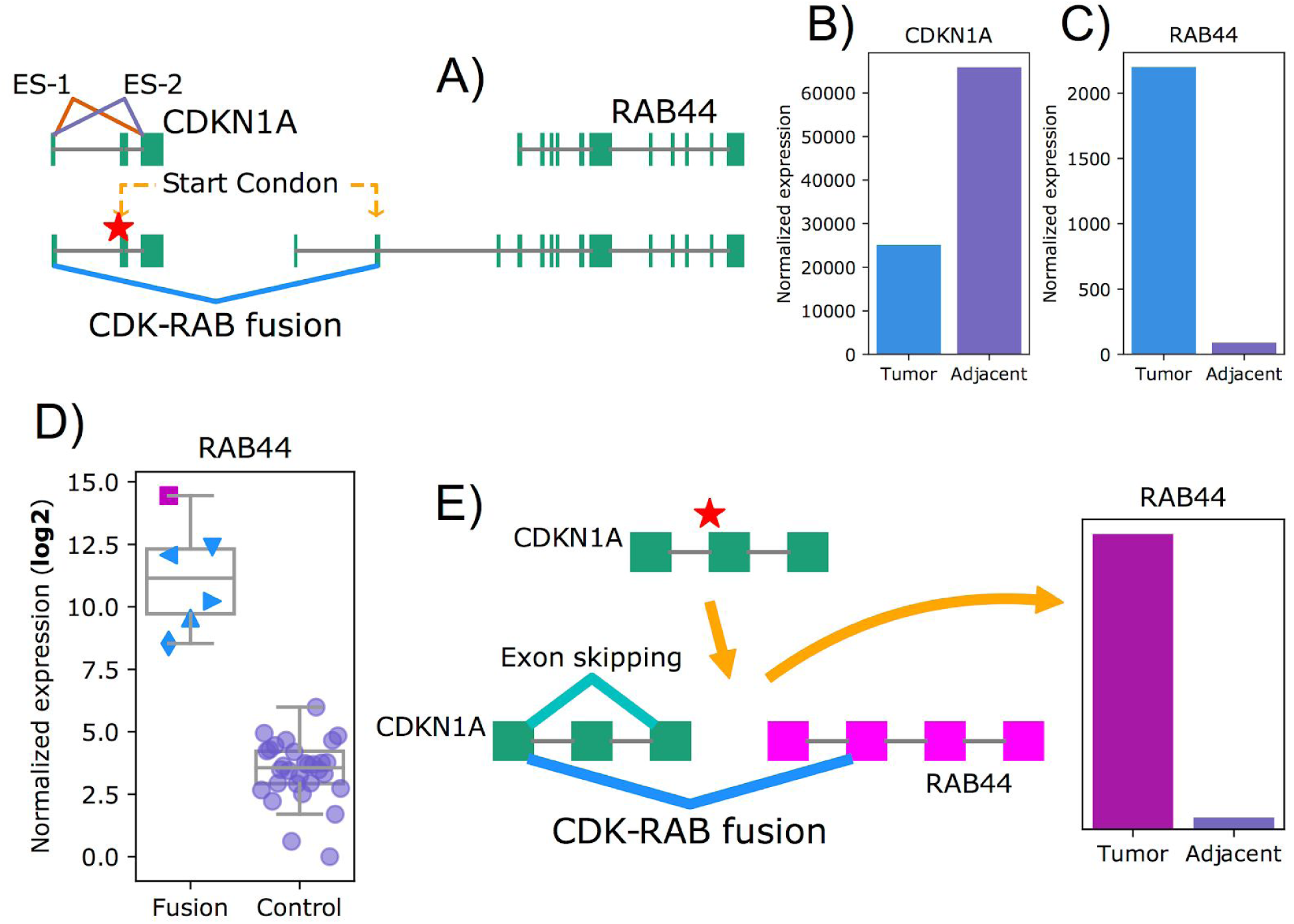
Transcript fusion between *CDKN1A* and *RAB44*, due to disruption of the splice site, activated *RAB44* expression. **(A)** Splice site somatic mutation associated with the exon skipping events within *CDKN1A* as well as the transcript fusion between *CDKN1A* and its downstream (1kb) gene *RAB44*. Two exon skipping events (ES-1 and ES-2) were shown because the boundary of the two isoforms of *CDKN1A* were slightly different but they were both observed in the RNA-Seq data. For the fusion, the second exon of *RAB44* was joined to the first exon of *CDKN1A*, which means the protein sequence of *RAB44* was joined to the UTR region of *CDKN1A* directly as both start codons of the two genes located in their second exons. **(B)** *CDKN1A* was downregulated in tumor when compared to tumor adjacent. **(C)** *RAB44* was upregulated. **(D)** All the six tumors, with mutations around the splice site, have higher *RAB44* expression when compared to randomly chosen tumor samples. Legends for the six tumors are in Supplementary Figure 3C. The magenta rectangle labelled patient, which has extremely high expression of *RAB44*, was discussed further in *TP53* pathway section. **(E)** Our model to explain these observations that one somatic mutation around the splice site is responsible for both the exon skipping and transcript fusion events. The fusion further leads to the activation of *RAB44*.

### CRISPR editing established the causality from splice site disruptive mutations to the transcript fusion and activation

To validate our model and establish the causality from the splice site mutations to the transcript fusion and activation, we knocked in mutations around the splice site of the second exon of *CDKN1A* in a lung cancer cell line, A549, using CRISPR/Cas9 (Supplementary Figure 5A). After confirming the presence of the mutations in clonally expanded cells with Sanger sequencing (Supplementary Figure 5B), we observed clearly the appearance of the fusion transcript in the edited cells rather than controls with a RT-PCR primer targeting the first exon of *CDKN1A* and the second exon of *RAB44* simultaneously (Figure 3A, Supplementary Figure 5C). RNA-Seq of three edited and three control cells further confirmed about the fusion transcript as well as the activation of *RAB44* expression (Figure 3B and 3C).

**Figure 3.**
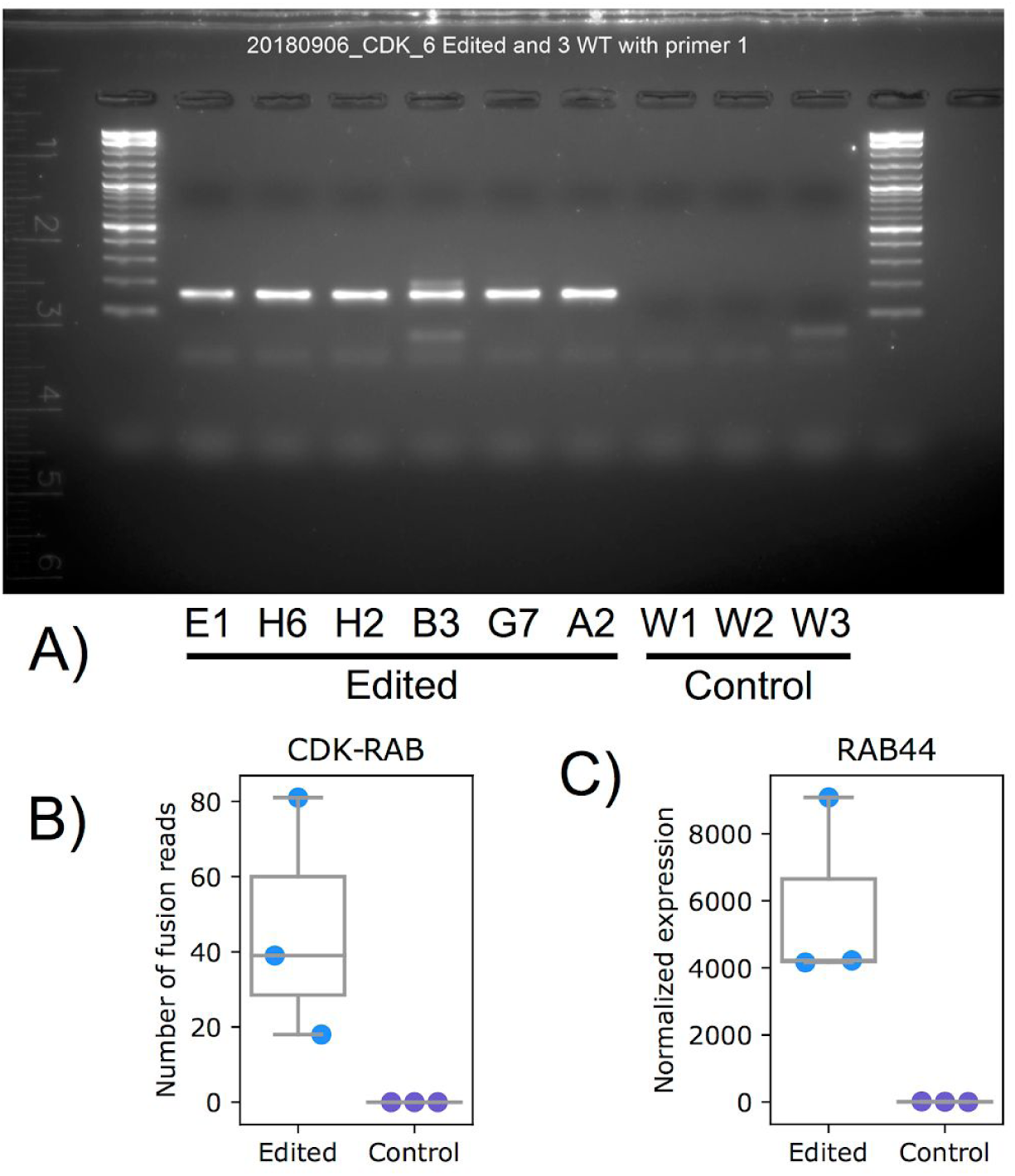
CRISPR editing established the causality from the splice site mutations to the transcript fusion and activation. **(A)** RT-PCR with a primer targeting both the first exon of *CDKN1A* and the second exon of *RAB44* verified the appearance of the fusion transcript in the edited cells, with merely the splice mutations knocked in, rather than in the control cells. **(B)** RNA-Seq reads further confirmed the joint between the first exon of *CDKN1A* and the second exon of *RAB44* directly on transcript level. **(C)** Upregulation of *RAB44* was clearly verified in RNA-Seq.

### Transcript fusion escapes exosome degradation of the readthrough transcripts

Fusion transcript appeared after knockin of merely splice site mutations. One narrative question is how these two independent genes could be joined together on the transcriptional level. A simple model is there is a readthrough transcript between these two genes even in normal cells which don’t carry such splice site mutations, however, it couldn’t be observed usually because it is degraded efficiently. On the other side, the fusion transcript, which undergoes proper splicing on the readthrough transcript, could be able to escape from the degradation and thus be visible in the mutated cells (Supplementary Figure 6A). Inhibiting degradation pathways is a rational way to check about this model. First of all, the nonsense-mediated decay (NMD) pathway, which degrades transcripts from 5’ to 3’, was less likely involved here, given that we couldn’t see difference on the region of these two genes when knocking down a few key components of this pathway, including *UPF1*, *UPF2*, *SMG6*, and *SMG7*, as shown in Supplementary Figure 6B and 6C, based on the data from ENCODE project and a published study^23^. Next, published *EXOSC3* knockdown and nuclear fraction nascent sequencing data^24^ in HeLa cells suggested the involvement of exosome associated pathway, which degrades RNAs from 3’ to 5’, as well as the restriction of such readthrough transcripts within chromatin fraction (Supplementary Figure 6D and 6E). Furthermore, *EXOSC3* knockout sequencing data^25^ in mouse embryonic stem cells confirmed about the involvement of the exosome associated degradation as the signals were amplified significantly when *EXOSC3* was knocked out, given that the region of these two genes were conserved between human and mouse (Figure 4A and 4B).

**Figure 4.**
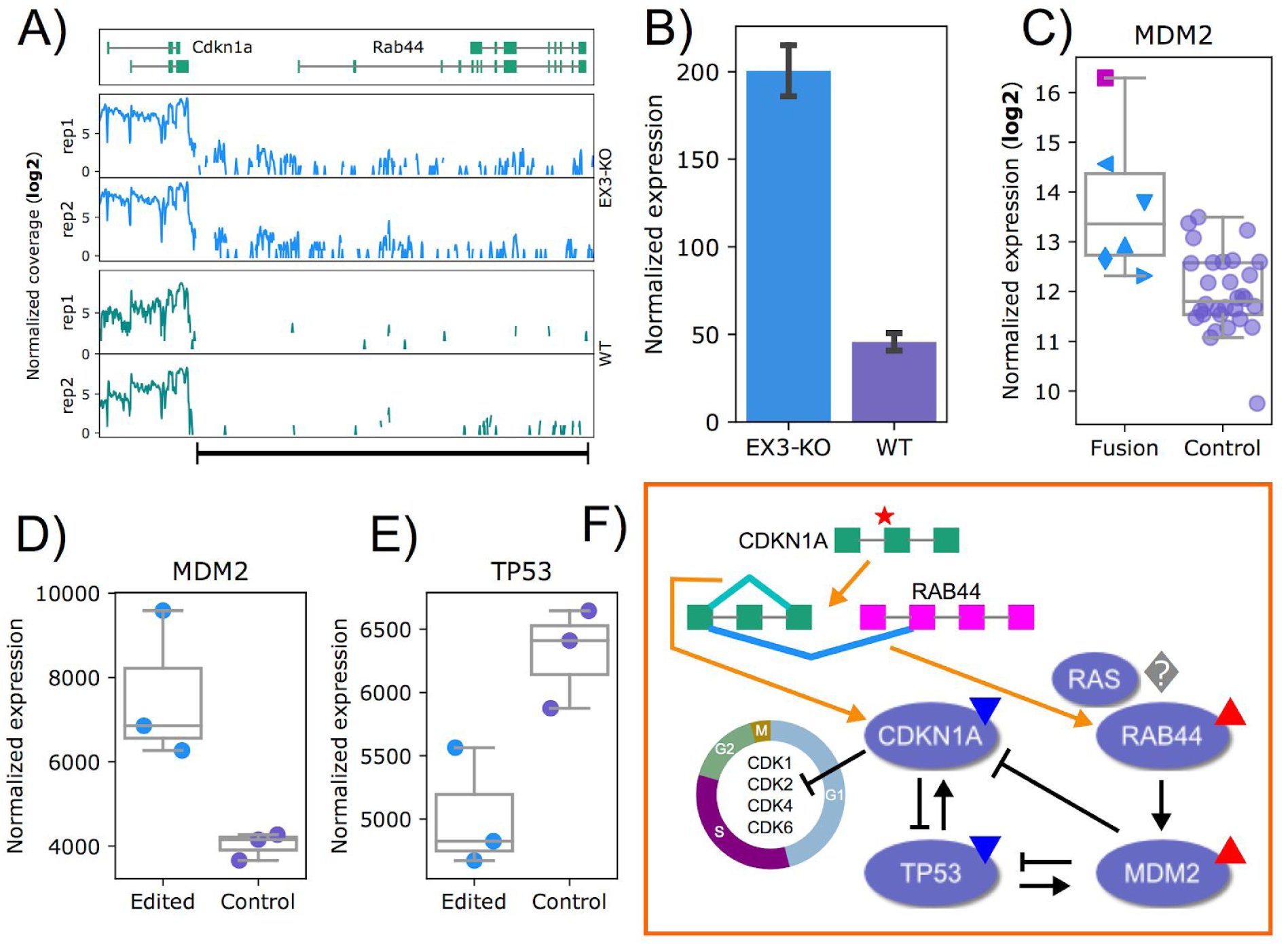
How the two genes could be possibly joined and what is the downstream effect. **(A)** Readthrough signals were amplified clearly in mouse embryonic stem cells after knocking out *EXOSC3*, a key component of the exosome complex. **(B)** Significant amplification of the readthrough signals on the regions indicated by the black bar line, including the intergenic region and the region of *RAB44*, in *EXOSC3* knocked out cells when compared to wide type cells, each with two biological replicates. Error bars were drawn with 95% confidence intervals. **(C)** A trend of upregulation of *MDM2* in those patients with the transcript fusion or the splice site mutations. Particularly, the magenta rectangle labelled patient had an extremely high expression of *MDM2* which correlated well with the extremely high expression of *RAB44* as shown in Figure 2D. The reason why this patient had such extreme expression of *RAB44* was further discussed in Supplementary Figure 7. Legends of these patients were in Supplementary Figure 3C. **(D)** Significant upregulation of *MDM2* was verified in the edited cells. **(E)** A trend of down regulation of *TP53* in edited cells. **(F)** Dual roles of the splice site mutation on the pathway perturbations. This mutation is responsible for the exon skipping events within *CDKN1A* that leads to downregulation of *CDKN1A* and benefits to cell growth. In the meanwhile, this mutation can also give rise to the transcript fusion which further leads to the upregulation of *RAB44*, upregulation of *MDM2*, inactivation of *TP53*, inactivation of *CDKN1A*, and finally acts on cell cycle and growth too. Blue lower triangular and red upper triangular indicates down and up regulation respectively.

### *RAB44* activation inactivates *TP53* pathway through upregulating *MDM2*

What is the downstream effect of the activation of *RAB44* in those cells which normally don’t express it? First of all, as shown in Figure 4C, cancer patient data implied the up-regulation of *MDM2*, which was a principle negative regulator of the p53 tumor suppressor. Particularly, *MDM2* was extremely up upregulated in a patient (TCGA-BT-A42C), which correlated very well with the *RAB44* expression as shown previously in Figure 2D. In fact, this patient had extreme expression of *RAB44* was less likely due to methodological artifacts or sampling noise but because the cells expressed a second type of fusion that joined the fourth exon of *RAB44* to the first region of *CDKN1A* (Supplementary Figure 7). Interestingly, there is a start codon in the fourth exon according to the second isoform of *RAB44*, thus this kind of fusion also doesn’t change its protein sequence. Furthermore, our in vitro experiments confirmed the significant up-regulation of *MDM2*, as well as the trend of down-regulation of *TP53*, in the edited cells as shown in Figure 4D and 4E. Besides, many genes, belonging to various cancer-related pathways, such as growth factor signaling and angiogenesis, were also deregulated (Supplementary Table 2 and 3).

Function of *RAB44* has not been well studied yet^26^, though it belongs to *RAS* oncogene family and has GTPase activity, which involves in membrane trafficking. However, even leaving aside of these roles, the processes here are still particularly interesting, considering that one mutation could give rise to exon skipping and transcript fusion simultaneously, while the skipping event further leads to down regulation of *CDKN1A*, which could activate multiple *CDKs* thus accelerate the cell growth, and the fusion event leads to up regulation of *RAB44*, up regulation of *MDM2*, inactivation of *TP53* and further down regulation of *CDKN1A* as summarized^27–31^ in Figure 4F.

## Discussion

We have already established the causality from the splice site disruptive mutations to the transcript fusion as well as the activation of *RAB44*. We also demonstrated one of the premises for such events to happen was the readthrough transcripts which occurred even in wild type cells although they might be degraded efficiently by the exosome complex. One more question is how the cells could recognize the readthrough transcripts while allowing those properly spliced fusion transcripts escape from degradation (Supplementary Figure 8A). Considering that *EXOSC3* was involved in degrading RNAs containing AU-rich elements (AREs)^32^, we checked one of the best-characterized motifs, AUUUA, in the region of these two genes. As shown in Supplementary Figure 8B, it was as expected that the frequency of this motif was higher in introns and the intergenic region than in exons, which suggested the readthrough transcripts, containing introns and the intergenic regions, might more easily be recognized and degraded by the complex.

It is essential to determine whether such splice site mutations or transcript fusions could be able to drive carcinogenesis. It is not easy to have a clear answer given that cancer initiation and development usually takes a long time and requires accumulation of additional mutations. As a preliminary attempt, we tried to check whether those six patients, with splice site mutations and transcript fusion events, carry any common somatic mutations. As shown in Supplementary Figure 9, *CDKN1A* is the only gene that mutated in all six patients, however, we can’t exclude the possibility of contributions from other genes, such as *BRCA2*, *FGFR3* or *EP300*, as they also mutated in multiple samples though they are less frequent and the mutations themselves are far away from each other, unlike those in *CDKN1A* that are all around the splice site.

The cell line we engineered to establish the causality was a lung cancer cell line (A549), which was different from those cancer tissues where we observed signals originally, including bladder, stomach and skin. This suggests the fusion and activation events are probably not cell type specific. Therefore, cancer sub-classification and genetic testing shouldn’t be limited in one or few cancer types. Besides of diagnosis, targeted therapy is worth further study, especially given that the fusion transcript or *RAB44* is only highly expressed in tumors rather than tumor adjacent tissues. Not mentioning that *RAB44* itself is not expressed in majority of human tissues except blood. Theoretically, even the drugs that target the transcript fusion or *RAB44* might not be easily developed in the near future, the diagnosis could still be valuable in clinical practice, considering that this group of patients might all undergo malfunction of *TP53*. Finally, our finding also warrants further attentions about the compound effects on transcription level between splicing disruptive mutations, including those on splice sites or splicing elements, and readthrough events, which seems to be more pervasive under cellular stress ^33^, that could lead to activation of oncogenes or inactivation of tumor suppressors through deregulating gene expression or giving rise to novel proteins.

## Methods

### Analysis of sequencing data from cancer patients and cell lines

Whole exome sequencing and RNA-Seq data of both tumor and tumor-adjacent samples of 657 patients in TCGA project, including 23 types of cancers, were downloaded from GDC database through application to dbGaP^34^. In addition, the same type of data in three published studies^16–18^, including 68 colon cancer, 22 small cell lung cancer, and 32 gastric cancer, was accessed from the European Genome-phenome Archive^35^ with approval from the data access committee. First of all, somatic mutations called from whole exome sequencing data were simply accessed either from the MAF files provided by GDC, including the results from both MuTect2 and VarScan2, or from the supplementary tables of each study with the coordinates converted from GRCh37 to GRCh38 to be consistent with the TCGA dataset. Next, on the RNA-Seq data, we calculated both inclusive reads (IR) and exclusive reads (ER) for every exon in each tumor and tumor-adjacent, as illustrated in Supplementary Figure 1B. Those skipping exons, supported by exclusive reads, when occurred only in tumors instead of tumor adjacent were of particular interest, given that they might be explained by cis effect from somatic mutations nearby. To confidentially detect such exon skipping events, we implemented a basic Bayesian hierarchical model, which takes advantage of shrinkage through partial pooling information across exons and samples (Supplementary Figure 1A). After approximating posterior distributions using automatic differentiation variational inference (ADVI) in PyMC3 (Supplementary Figure 1C), we estimated the difference of percent spliced-in index (ΔPSI, PSI = IR / (IR + ER)) for each exon between tumor and tumor-adjacent. Confident events were those exons with highest posterior density (HPD, alpha=95%) of ΔPSI excluding the value of zero. We reported the association candidates for these events if somatic mutations were identified either deeply within the exon, for exonic splicing enhancers (ESEs), or around the splice site (± 20bp). 20bp was determined according to the parameters used by MaxEntScan^19^. To find independent evidence for these candidates, we further checked additional tumor samples from the TCGA project, without matched tumor adjacent samples sequenced, as well as 935 cancer cell lines from the CCLE project^36^.

### In vitro CRISPR/Cas9 editing of A549 cell line

Before becoming our choice of cell line, RNA-Seq data of A549 from CCLE project was examined to be sure that it didn’t carry the fusion transcript, didn’t express *RAB44* and didn’t have disruptive mutations around the splice site of the second exon of CDKN1A.

> **gRNA:**CTTCCTTGTATCTCTGCTGC<u>AGG</u>
>
> **RT-PCR Primer:** GCCGAAGTCAGTTCCTTGTG (+, within exon 1 of CDKN1A) and ACCATCAGCTGGCTCTCTTG (-, within exon 2 of RAB44)

Libraries for three edited samples and three wild type controls were processed following TruSeq RNA library prep kit v2 protocol (Illumina). RNA-Seq reads were aligned to GRCh38, with the gene annotation file downloaded from ENSEMBL (v94), using STAR (v2.6.1c). Gene expression matrix was collected using featureCounts (v1.6.2). Differential expression analysis was performed with DESeq2 (v1.20.0). Raw sequencing data were deposited in SRA database under accession PRJNA513099.

### Evidence for exosome degradation of readthrough transcripts

Fastq files of these two studies^24, 25^ were downloaded from GEO (GSE81662) and SRA (SRP042355) and then aligned to GRCh38 and GRCm38 using STAR respectively. Coverage signal was calculated on the bedGraph files converted from bam files using bamCoverage (deepTools, v3.1.3) and further normalized by median-like size factors determined by DESeq2.

## Author Contributions

H.S., W.W., L.M.S. designed the project, H.S. analyzed the data, Z.C., H.Z., W.M. did CRISPR/Cas9 editing, M.N., K.R., P.J. performed sequencing and other experiments, H.S., L.M.S., W.W., C.Z., J.W., R.A. prepared the manuscript.

## Acknowledgements

We would like to thank Lars Velten, Shendi Li, Derek Anolles, Yuehan Feng, Michael Sikora, Fan Zhou, Binqing Zhao and Ying Liu for helpful discussion and calm support. We would like to thank TCGA Research Network and European Genome-phenome Archive (EGA) for sharing with us raw sequencing data of the cancer cohorts. We also would like to thank all those patients donating biopsy for supporting the cancer research.

## Reference

1. Dvinge, H., Kim, E., Abdel-Wahab, O. & Bradley, R. K. RNA splicing factors as oncoproteins and tumour suppressors. Nat. Rev. Cancer 16, 413–430 (2016).

2. Scotti, M. M. & Swanson, M. S. RNA mis-splicing in disease. Nat. Rev. Genet. 17, 19–32 (2015).

3. Cieply, B. & Carstens, R. P. Functional roles of alternative splicing factors in human disease. Wiley Interdiscip. Rev. RNA 6, 311–326 (2015).

4. Guo, W. et al. RBM20, a gene for hereditary cardiomyopathy, regulates titin splicing. Nat. Med. 18, 766–773 (2012).

5. Vaz-Drago, R., Custódio, N. & Carmo-Fonseca, M. Deep intronic mutations and human disease. Hum. Genet. 136, 1093–1111 (2017).

6. Jayasinghe, R. G. et al. Systematic Analysis of Splice-Site-Creating Mutations in Cancer. Cell Rep. 23, 270–281.e3 (2018).

7. Fairclough, R. J., Wood, M. J. & Davies, K. E. Therapy for Duchenne muscular dystrophy: renewed optimism from genetic approaches. Nat. Rev. Genet. 14, 373–378 (2013).

8. Reungwetwattana, T., Liang, Y., Zhu, V. & Ou, S.-H. I. The race to target MET exon 14 skipping alterations in non-small cell lung cancer: The Why, the How, the Who, the Unknown, and the Inevitable. Lung Cancer 103, 27–37 (2017).

9. Lee, S. C.-W. & Abdel-Wahab, O. Therapeutic targeting of splicing in cancer. Nat. Med. 22, 976–986 (2016).

10. Mou, H. et al. CRISPR/Cas9-mediated genome editing induces exon skipping by alternative splicing or exon deletion. Genome Biol. 18, 108 (2017).

11. Roy, K. R. et al. Multiplexed precision genome editing with trackable genomic barcodes in yeast. Nat. Biotechnol. 36, 512–520 (2018).

12. Findlay, G. M., Boyle, E. A., Hause, R. J., Klein, J. C. & Shendure, J. Saturation editing of genomic regions by multiplex homology-directed repair. Nature 513, 120–123 (2014).

13. Findlay, G. M. et al. Accurate classification of BRCA1 variants with saturation genome editing. Nature 562, 217–222 (2018).

14. Li, H., Wang, J., Mor, G. & Sklar, J. A Neoplastic Gene Fusion Mimics Trans-Splicing of RNAs in Normal Human Cells. Science 321, 1357–1361 (2008).

15. Rickman, D. S. et al. SLC45A3-ELK4 is a novel and frequent erythroblast transformation-specific fusion transcript in prostate cancer. Cancer Res. 69, 2734–2738 (2009).

16. Seshagiri, S. et al. Recurrent R-spondin fusions in colon cancer. Nature 488, 660–664 (2012).

17. Liu, J. et al. Integrated exome and transcriptome sequencing reveals ZAK isoform usage in gastric cancer. Nat. Commun. 5, 3830 (2014).

18. Rudin, C. M. et al. Comprehensive genomic analysis identifies SOX2 as a frequently amplified gene in small-cell lung cancer. Nat. Genet. 44, 1111–1116 (2012).

19. Yeo, G. & Burge, C. B. Maximum entropy modeling of short sequence motifs with applications to RNA splicing signals. J. Comput. Biol. 11, 377–394 (2004).

20. Crooks, G. E., Hon, G., Chandonia, J.-M. & Brenner, S. E. WebLogo: a sequence logo generator. Genome Res. 14, 1188–1190 (2004).

21. McVety, S. Disruption of an exon splicing enhancer in exon 3 of MLH1 is the cause of HNPCC in a Quebec family. J. Med. Genet. 43, 153–156 (2005).

22. Korvatska, O. et al. Altered splicing of ATP6AP2 causes X-linked parkinsonism with spasticity (XPDS). Hum. Mol. Genet. 22, 3259–3268 (2013).

23. Colombo, M., Karousis, E. D., Bourquin, J., Bruggmann, R. & Mühlemann, O. Transcriptome-wide identification of NMD-targeted human mRNAs reveals extensive redundancy between SMG6- and SMG7-mediated degradation pathways. RNA 23, 189–201 (2017).

24. Schlackow, M. et al. Distinctive Patterns of Transcription and RNA Processing for Human lincRNAs. Mol. Cell 65, 25–38 (2017).

25. Pefanis, E. et al. RNA exosome-regulated long non-coding RNA transcription controls super-enhancer activity. Cell 161, 774–789 (2015).

26. Yamaguchi, Y. et al. Rab44, a novel large Rab GTPase, negatively regulates osteoclast differentiation by modulating intracellular calcium levels followed by NFATc1 activation. Cell. Mol. Life Sci. 75, 33–48 (2018).

27. El-Deiry, W. WAF1, a potential mediator of p53 tumor suppression. Cell 75, 817–825 (1993).

28. Zhang, Z. et al. MDM2 Is a Negative Regulator of p21WAF1/CIP1, Independent of p53. J. Biol. Chem. 279, 16000–16006 (2004).

29. Broude, E. V. et al. p21 (CDKN1A) is a Negative Regulator of p53 Stability. Cell Cycle 6, 1467–1470 (2007).

30. Shi, D. & Gu, W. Dual Roles of MDM2 in the Regulation of p53: Ubiquitination Dependent and Ubiquitination Independent Mechanisms of MDM2 Repression of p53 Activity. Genes Cancer 3, 240–248 (2012).

31. Wu, X., Bayle, J. H., Olson, D. & Levine, A. J. The p53-mdm-2 autoregulatory feedback loop. Genes Dev. 7, 1126–1132 (1993).

32. Mukherjee, D. et al. The mammalian exosome mediates the efficient degradation of mRNAs that contain AU-rich elements. EMBO J. 21, 165–174 (2002).

33. Vilborg, A. & Steitz, J. A. Readthrough transcription: How are DoGs made and what do they do? RNA Biol. 14, 632–636 (2017).

34. Tryka, K. A. et al. NCBI’s Database of Genotypes and Phenotypes: dbGaP. Nucleic Acids Res. 42, D975–9 (2014).

35. Lappalainen, I. et al. The European Genome-phenome Archive of human data consented for biomedical research. Nat. Genet. 47, 692–695 (2015).

36. Barretina, J. et al. The Cancer Cell Line Encyclopedia enables predictive modelling of anticancer drug sensitivity. Nature 483, 603–607 (2012).

